# A widely distributed genus of soil Acidobacteria genomically enriched in biosynthetic gene clusters

**DOI:** 10.1101/2021.05.10.443473

**Authors:** Alexander Crits-Christoph, Spencer Diamond, Basem Al-Shayeb, Luis Valentin-Alvarado, Jillian F. Banfield

## Abstract

Bacteria of the phylum Acidobacteria are one of the most abundant bacterial across soil ecosystems, yet they are represented by comparatively few sequenced genomes, leaving gaps in our understanding of their metabolic diversity. Recently, genomes of Acidobacteria species with unusually large repertoires of biosynthetic gene clusters (BGCs) were reconstructed from grassland soil metagenomes, but the degree to which these species are widespread is still unknown. To investigate this, we augmented a dataset of publicly available Acidobacteria genomes with 46 metagenome-assembled genomes recovered from permanently saturated organic-rich soils of a vernal (spring) pool ecosystem in Northern California. We recovered high quality genomes for three novel species from *Candidatus* Angelobacter (a proposed subdivision 1 Acidobacterial genus), a genus that is genomically enriched in genes for specialized metabolite biosynthesis. Acidobacteria were particularly abundant in the vernal pool sediments, and a *Ca*. Angelobacter species was the most abundant bacterial species detected in some samples. We identified numerous diverse biosynthetic gene clusters in these genomes, and also in additional genomes from other publicly available soil metagenomes for other related *Ca*. Angelobacter species. Metabolic analysis indicates that *Ca*. Angelobacter likely are aerobes that ferment organic carbon, with potential to contribute to carbon compound turnover in soils. Using metatranscriptomics, we identified in situ expression of specialized metabolic traits for two species from this genus. In conclusion, we expand genomic sampling of the uncultivated *Ca*. Angelobacter, and show that they represent common and sometimes highly abundant members of dry and saturated soil communities, with a high degree of capacity for synthesis of diverse specialized metabolites.

## Introduction

It is estimated that an overwhelming majority of soil bacterial species have thus far been recalcitrant to cultivation (Steen et al. 2019), and these uncultivated bacteria are not evenly distributed across the tree of life (Lloyd et al. 2018). While many phyla of bacteria found in soils have few cultivated representatives, there are few as ubiquitous and diverse as the Acidobacteria (A. M. Kielak et al. 2016). From metagenomic and 16S rRNA surveys we have learned that Acidobacteria are collectively the most abundant phylum in soils (Fierer 2017), harboring significant taxonomic diversity (Lee, Ka, and Cho 2008; A. Kielak et al. 2009), with over 26 accepted subdivisions (Barns et al. 2007). However, they are relatively undersampled in cultivation efforts, with fewer than 100 sequenced isolate genomes from the entire phylum deposited into the RefSeq database as of the start of 2021. Some reported isolates have also not been genomically sequenced or deposited into public strain collections, complicating the study of even previously cultivated members of the phylum. Isolate-based studies of soil Acidobacteria indicate that they are often heterotrophic, aerobic and capable of complex carbon degradation, and are thought to be mostly oligotrophic (A. M. Kielak et al. 2016). However, it is unclear to what degree these findings extrapolate to the diversity of the entire phylum in soils.

More recently, genome-resolved metagenomics, or the process of assembling and curating genomes directly from metagenomes, has been applied to soil bacterial communities and resulted in the assembly of hundreds of novel soil Acidobacteria genomes (Diamond et al. 2019); (Woodcroft et al. 2018); (Woodcroft et al. 2018; Xue et al. 2020). Genome annotation and functional prediction from these genomes have improved our understanding of the phylum’s metabolic potential. Metabolic characteristics predicted from genomes of uncultivated Acidobacteria include the ability to degrade many complex carbohydrate compounds via carbohydrate active enzymes, the capacity for nitric oxide reduction, and the ability to oxidize methanol (Diamond et al. 2019). It also was previously shown that many Acidobacteria genomes from metagenomic data from a single soil ecosystem encode numerous gene clusters for the biosynthesis of specialized metabolites (Crits-Christoph et al. 2018). In particular, two lineages of Acidobacteria in subgroup 1 and 4, designated *Candidatus* Angelobacter (Genome Taxonomy Database genus g__Gp1-AA17; NCBI taxon ‘Acidobacteria bacterium AA117’) and *Candidatus* Eelbacter (NCBI taxonomy Blastocatellia bacterium AA13), respectively, were found to possess large repertoires of biosynthetic genes. Specifically, the genomes each encoded for 300-400 Kb of nonribosomal peptides synthetases (NRPSs) and polyketides synthases (PKSs). A third lineage of uncultivated Acidobacteria has also been reported which also possessed similarly large NRPS and PKS gene clusters, and was sequenced from ocean biofilm samples (W. Zhang et al. 2019). Additional genomes from this third lineage have since been noted for their unusually large nonribosomal peptide genes (Nayfach et al. 2020). However, from the genomes reported thus far, it remains unclear how these three lineages are related, or whether they are widespread in soil environments. Here, we extend these results using publicly available Acidobacteria genomes from a variety of soil types, and augmented the dataset by additional sampling and new metagenomic analysis of saturated soils from a vernal pool ecosystem. Of the previously reported Acidobacteria lineages enriched in biosynthetic gene clusters, we identified several more species from *Ca*. Angelobacter, and focused our analysis on this group. Our results substantially increase the number of reported genomes from the *Ca*. Angelobacter genus and suggest that a significant investment in secondary metabolism is a common feature of soil bacteria from this lineage.

## Methods

### Field sampling

We collected soil samples from a seasonal vernal pool in Lake County, California, in October 2018 and October 2019. Samples were frozen in the field using dry ice, and kept at −80 C until extraction. The Qiagen PowerSoil Max DNA extraction kit was used to extract DNA from 10 g of soil, and the Qiagen AllPrep DNA/RNA extraction kit was used to extract RNA from 2 g of soil. Samples were sequenced by the QB3 sequencing facility at the University of California, Berkeley on a NovaSeq 6000. Read lengths for the 2018 DNA samples and the RNA samples were 2×150 bp and 2×250 bp for the 2019 DNA samples. A sequencing depth of 10 Gb was targeted for each of the 2018 samples, and 20 Gbp for each of the 2019 samples. Geochemical measurements (concentrations of Total Carbon, Total Nitrogen, Calcium, Zinc, Magnesium, Copper, and Iron) were performed on 10 selected soil samples from the sampling site at the University of California, Davis Analytical Laboratory (https://anlab.ucdavis.edu/methods-of-analysis) using publicly available protocols.

### Metagenomic assembly and annotation

Metagenomic sequencing reads were assembled using the IDBA_UD assembler (Peng et al. 2012). Contigs greater than 2.5 Kb were retained and sequencing reads from all samples were cross-mapped against each resulting assembly using Bowtie2 (Langmead and Salzberg 2012). The resulting differential coverage profiles were filtered at a 95% read identity cutoff, and then used for genome binning with MetaBAT2 (Kang et al. 2019). Resulting genome bins were assessed for completeness and contamination using CheckM (Parks et al. 2015), and were manually curated using taxonomic profiling with GGKBase. Taxonomy was assigned to genome bins and a phylogenetic tree was constructed using phylogenetic placement of single copy marker genes with GTDB-Tk (Chaumeil et al. 2019). Angelobacter genomes were identified by classification with GTDB-Tk as either family ‘Gp1-AA117’ or genus ‘Gp1-AA17; nomenclature derived from the first identified genome Angelobacter Gp1-AA117. Community relative abundance profiles were determined for each sample using the GraftM (Boyd, Woodcroft, and Tyson 2018) metagenomic classifier and the ribosomal protein L6 marker gene.

Biosynthetic gene clusters were annotated in genomes using antiSMASH 5.0 (Blin et al. 2019). The number of Condensation domains and Ketoacyl synthase domains was determined by using HMMER3 (Eddy 2011) and querying all antiSMASH predicted biosynthetic proteins using the PF00109 and PF00668 Pfam HMMs. BiG-SCAPE (Navarro-Muñoz et al. 2020) was used to generate gene cluster families of BGCs. KEGG functional annotations for genes across the entire genomes were obtained using METABOLIC (Zhou et al. 2020).

Metatranscriptomic data was mapped to all genome bins using Bowtie2, and filtered to only paired end reads with >95% identity using Python. Read counts were then log-normalized in R.

## Results

### Sampling and metagenomic sequencing

We collected 29 soil samples from a seasonal vernal pool in Lake County, California, USA in October 2018 and October 2019 (**Supplementary Table S1**). The elevation of the site is approximately 600 m and the pool is surrounded by Douglas Fir and Oak (**Fig 1b**). The soils at the site are fine grained, organic-rich mud and clay-rich at depth, surrounded by gravelly loam Inceptisols formed in material weathered from rhyolitic tuff. Sampling occurred when the pool was at its driest before the first major autumn rainfall, along a transect in the pool bed that would be covered by water for a majority of the year. Total nitrogen and total carbon measured at the site averaged 1% and 13% respectively, both decreased with increasing soil depth (**Supplementary Table S2**). The soils were slightly less carbon rich at depth, where by 80 cm total carbon decreased to <10%. All samples were saturated with water at the time of collection. Samples were collected from 25, 35, 45, 60, and 80 cm depths, and either stored on dry ice for DNA extraction or flash frozen in ethanol cooled with dry ice for RNA extraction.

**Figure 1:**
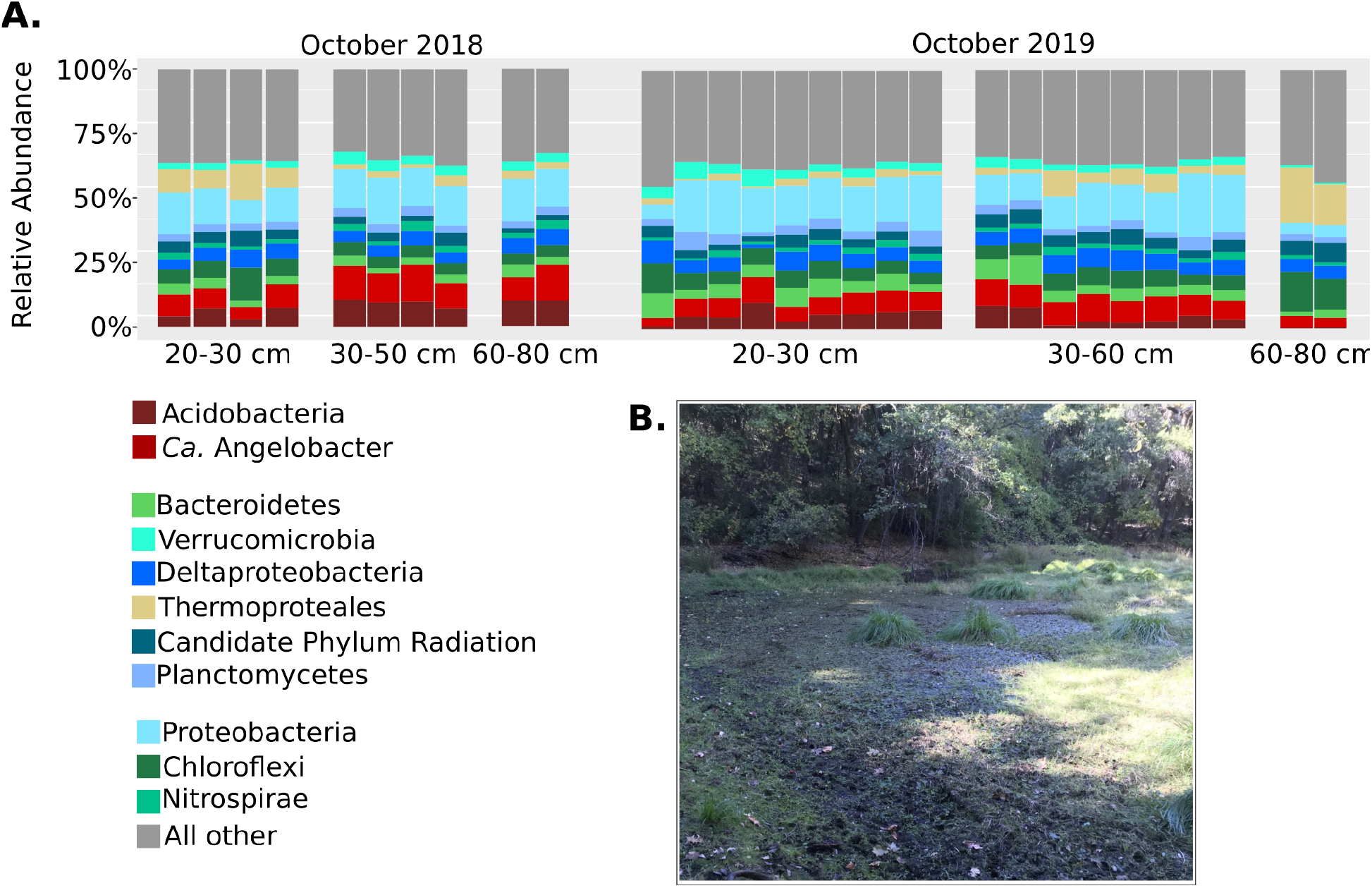
Community composition of vernal pool soils. (**A**) Ribosomal protein (L6) abundances and taxonomic classifications across all metagenomic samples obtained in this study. The abundances of *Ca*. Angelobacter (*Gp1-AA117)* are shown separately from all other hits in phylum Acidobacteria. (**B**) Photograph of the vernal pool that was metagenomically sampled in this study, in Lake County, California, USA.

Sample metagenomes were sequenced to an average depth of 10 Gb per sample for 2018 and 20 Gb per sample for samples collected in 2019. Metagenomic assembly resulted in on average 479 Mb of assembled sequence in contigs > 1 Kb per sample. Using these assemblies, we generated 46 dereplicated Acidobacterial genomes of at least medium-quality (>90% complete, <10% contaminated) (**Supplementary Table S3**).

### Community composition and assembled genomes

In order to assess the bacterial community composition of the vernal pool, the L6 ribosomal protein was used as a phylogenetic marker with the GraftM software (Boyd, Woodcroft, and Tyson 2018). The bacterial communities at the site were found to be less complex than the soil communities observed by a previous metagenomic effort in a arid grassland meadow (Diamond et al. 2019). Dominant taxa included species in the phylum Acidobacteria, Proteobacteria, Chloroflexi, with a variety of Archaea dominating the community at deeper depths of 60-80 cm (**Fig 1a**). At high taxonomic ranks of bacteria and archaea, community composition was fairly consistent within each stratum across depths. Members of the Candidate phyla radiation consistently composed ~10% of the community, more than has been previously reported in drier soils (Sharrar et al. 2020; Nicolas et al. 2020).

Of particular interest was the high abundance of the phylum Acidobacteria within the vernal pool soil microbial community, in some samples reaching 25% relative abundance. The 46 assembled near-complete species-dereplicated Acidobacteria genomes from the site were placed in a phylogenetic tree of all 370 known Acidobacteria species in the NCBI Assembly database and the Genome Taxonomy Database (GTDB) using a concatenated set of ribosomal proteins (**Fig 2**). Genomes recovered from the vernal pools samples derived from five different Acidobacteria classes (group 1, 3, 5, 7, and 11), indicating a wide diversity of species abundant for this phylum in the vernal pool.

**Figure 2:**
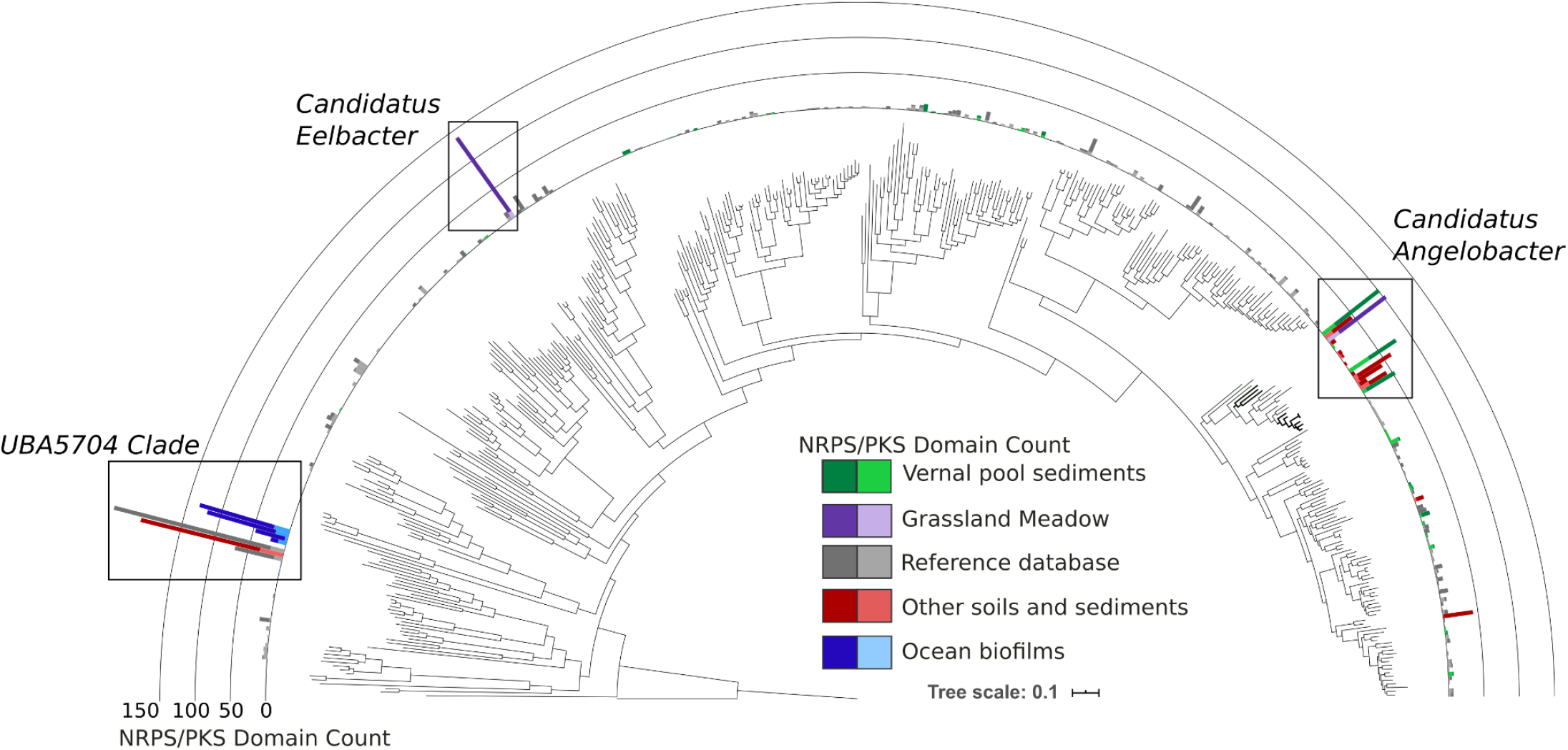
Phylogeny of the Acidobacteria annotated with biosynthetic domain counts per genome. Concatenated ribosomal protein phylogeny of all Acidobacteria genomes in NCBI GenBank, additional genomes obtained from IMG, and genomes obtained from this study (“Vernal pool sediments”). Plotted is the number of biosynthetic NRPS CD /PKS KS domains per genome, and genomes are colored by their ecosystem of origin.

Phylogenetic analysis of single copy marker genes identified three novel genomes related to the uncultivated group 1 Acidobacteria, *Ca*. Angelobacter (GTDB genus g__Gp1-AA17; NCBI taxon ‘Acidobacteria bacterium AA117’). Besides the previously published genome from another site in Northern California, there was only one other metagenome assembled genome from this clade, recovered from a thawing permafrost peatland located in arctic Sweden. The three new genomes obtained from the vernal pool study site were all near-complete with low estimated contamination and ranged in size from 6-7 Mb, with an average GC content of 55% (**Supplementary Table S4**). One of these genomes, *SRVP-Angelobacter-2*, was 6.44 Mb total across 39 contigs, which is the most contiguous assembly of a *Candidatus Angelobacter* species, and significantly more contiguous than the previously published genome. Assembly of large contiguous genome fragments is important for accurate binning and especially for the recovery of complete BGCs and identification of nearby genomic features. *Angelobacter* were at reasonably high abundances in all samples, and the *SRVP-Angelobacter-3* species was the most abundant organism in samples from 20 cm depth. Two genomes contained a complete 16S rRNA gene with 95.2% sequence identity, consistent with being members of the same genus.

To further expand our characterization of the *Ca*. Angelobacter genus, we searched the IMG database of assembled metagenome bins for additional genomes from the genus by phylogenetic placement of all acidobacterial genomes in the dataset. We identified additional draft genomes for five more species in the *Ca*. Angelobacter, all from soils. Three were from a large metagenomic study of corn and switchgrass rhizosphere in Michigan (B. Zhang et al. 2017), one from a metagenomic study of soils amended with Pyrogenic organic matter in New York (Whitman et al. 2016), and one genome was obtained from a mini-metagenomic selection approach from Massachusetts forest soils (Alteio et al. 2020) (**Supplementary Table S4**). Comparing these genomes by average nucleotide identity (ANI), we found that six of the Angleobacter genomes clustered together with >60% ANI (**Fig S1**). The genome obtained from the New York study intriguingly shared 96.8% ANI with one of the genomes obtained from Michigan soils, indicating that they could be considered the same species based on a 95% ANI definition of microbial species; all of the other genomes appeared to be separate species.

### Many species of *Candidatus* Angelobacter genus possess diverse biosynthetic gene clusters

The previous genome reported from *Ca*. Angelobacter was notable in its substantial genomic capacity for production of specialized metabolites, particularly via large biosynthetic gene clusters of nonribosomal peptide synthetases and polyketide synthases. To understand how consistent this trait is across this genus, we ran antiSMASH 5.0 on all of the recovered genomes to identify Biosynthetic Gene Clusters (BGCs) and predicted both polyketide keto-synthase (KS) and NRPS condensation (CD) protein domains across antiSMASH BGCs. Visualizing the number of these biosynthetic domains found per genome, it is clear that the *Ca*. Angelobacter genus stands out in the acidobacterial phylum (**Fig 2**). We also phylogenetically place two other independent acidobacterial clades with large numbers of biosynthetic enzymatic domains. *Candidatus Eelbacter* is a group 4 Acidobacteria genome previously reported; there is also a lineage represented by genomes previously reported from ocean biofilm and soil metagenomes that cluster phylogenetically with each other and are identified by similarity to a genome deposited with the moniker ‘UBA5704’ (Parks et al. 2017). Within the *Ca*. Angelobacter clade, we also identified a minority of genomes with few biosynthetic gene clusters, indicating that genomically encoded biosynthetic capacity may vary within this lineage. This can also be the case for some members of the Actinomycetales, which are renowned for specialized metabolite production in general, yet the trait can be patchy across individual species (Parks et al. 2017; Chase et al., 2021). It is also possible that BGCs were not assembled or binned properly in some *Ca*. Angelobacter genomes from metagenomes.

The total number of BGCs in newly recovered *Ca*. Angelobacter genomes rivals, and in many cases outnumbers, the previously reported *Ca*. Angelobacter genome’s biosynthetic gene content (**Fig 3A**; **Supplementary Table S5**). Many *Ca*. Angelobacter BGCs were NRPS or NRPS-PKS hybrids with unusual complexity and length, and the NRPS genes within the clusters were unusually large, with the largest ORF in the genus being 24 Kbp in length. No biosynthetic gene cluster contained adjacent known genomic markers of siderophore biosynthesis (such as TonB-dependent receptors or periplasmic binding proteins), but were often associated with adjacent genes for MacB tripartite efflux pumps.

**Figure 3:**
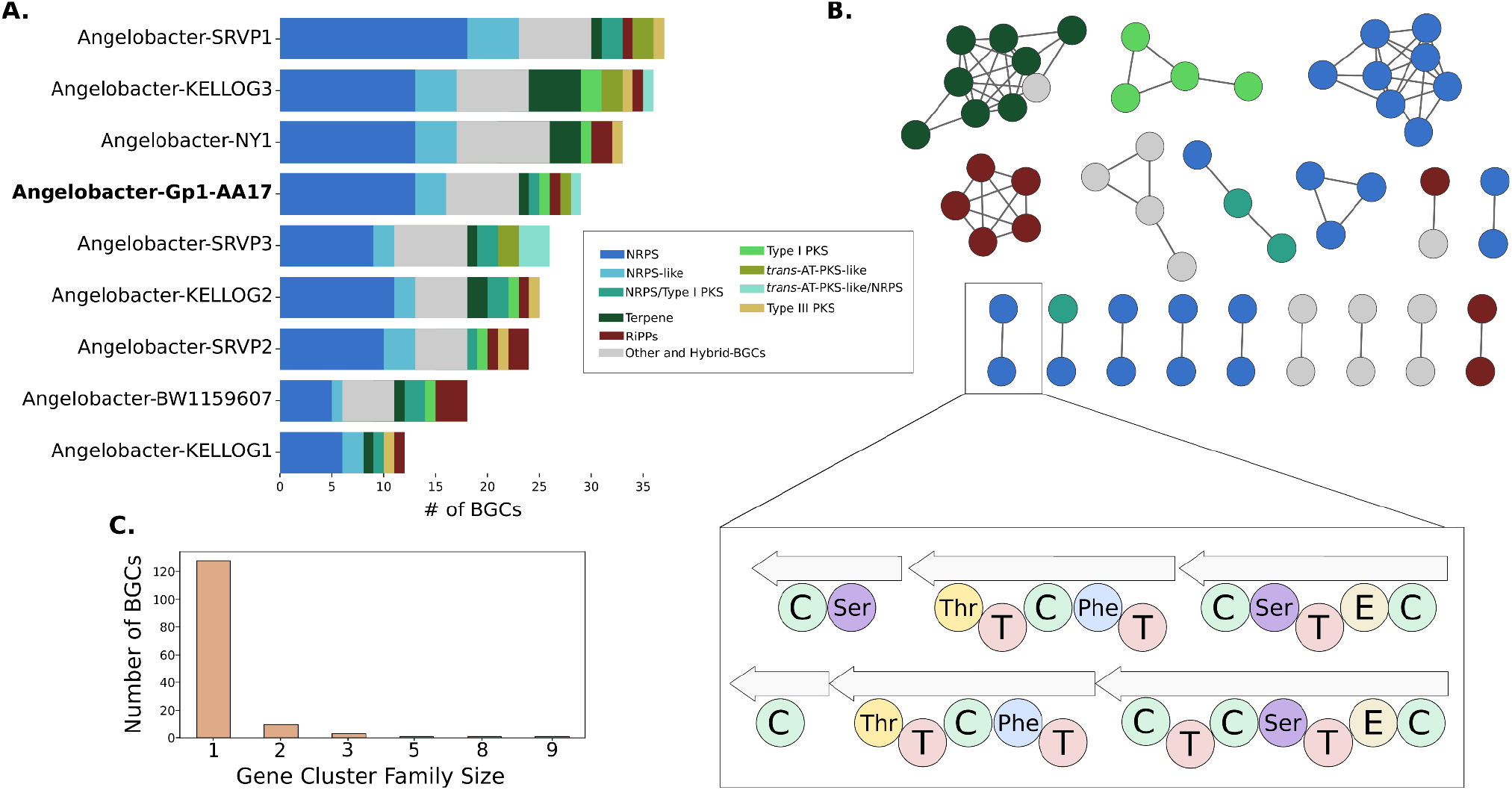
Biosynthetic gene clusters from genomes in *Candidatus* Angelobacter. (**A)** The number and class of BGCs in each species genome from *Ca*. Angelobacter. The previously published reference genome for this genus is highlighted in contrast to new genomes obtained in this study. (**B**) A BiG-SCAPE gene cluster family network of BGCs from *Ca*. Angelobacter. Each node is a BGC, connected to other similar BGCs by genomic similarity. Two core NRPS genomes from different species are shown in the inset. Nomenclature: C, Condensation Domain; Ser, Adenylation Domain (Serine); The, Adenylation Domain (Threonine); T, Peptidyl Carrier Protein; Phe, Adenylation Domain (Phenylalanine). E, Epimerase. (**C**) The number of *Ca*. Angelobacter BGCs in a gene cluster family of a certain size. The vast majority of BGCs are novel singletons.

To clarify whether *Ca*. Angelobacter species tend to share similar BGCs, we applied the BiG-SCAPE workflow to the recovered *Ca*. Angelobacter BGC collection to identify gene cluster families, or groups of related BGCs. We found that the majority of BGCs in the collection were singletons, indicating substantial genetic diversity and comparatively few BGCs shared between species (**Fig 3B, 3C**). Of BGC families that were shared by species, the majority were only shared by two species; only five clusters were shared amongst more than three *Ca*. Angelobacter species. The gene cluster families that were commonly shared include Terpene, Type I PKS, NRPS, and a ribosomally synthesized peptide gene cluster family. While most of the shared gene cluster families had fewer genes than average, intriguingly, two *Ca*. Angelobacter species from different study sites shared a highly similar large NRPS cluster with similar, but non-identical adenylation domain structure (**Fig 3B**).

Genomic inferences about primary metabolisms can help inform understanding of a microorganisms’ lifestyle and trophic niche, while also possibly guiding cultivation efforts. Generally the predicted metabolic capabilities of all 46 vernal pool acidobacteria identify them as aerobic heterotrophs with reasonable capacity for complex carbohydrate degradation and assimilation **(Fig S2**). Additionally, all of the genomes encode at least one (or more) of the following energy generation mechanisms that can operate when oxygen is not available: ethanol fermentation, dissimilatory nitrate reduction, or dissimilatory sulfate reduction. Given that these organisms encode a full set of respiratory complex enzymes, with an oxygen utilizing terminal oxidase, it is more likely that anaerobic metabolism is used in specific situations where oxygen is unavailable.

The large sizes of *Ca*. Angelobacter genomes are consistent with these organisms adopting a lifestyle strategy of metabolic versatility. We used the most complete and contiguous genome, *Angelobacter*-SRVP2, to further examine specific metabolic traits that allow for plasticity in carbon assimilation and energy generation. *Ca*. Angelobacter have complete pathways for glycolysis, oxidative pentose phosphate conversions, and the TCA cycle (**Supplementary Table S6**). They also encode a four complex oxygenic respiratory chain including an NADH:quinone oxidoreductase (Complex I), succinate dehydrogenase (Complex II), cytochrome b containing Complex III, and an oxygen utilizing cytochrome c oxidase (Complex IV). These genomes also encode a separate oxygen-utilizing cytochrome bd-like ubiquinol oxidase, which is thought to operate under lower oxygen availability (Poole and Cook 2000). Two putative mechanisms for anaerobic energy generation are also present: fermentation to ethanol and the ability to reduce nitrate to nitrite (**Supplementary Table S6**). They have the ability to assimilate acetate (and other 2 carbon compounds) into biomass by encoding enzymes for the glyoxylate shunt. This provides additional metabolic flexibility for situations where complex carbohydrates or hexose sugars are unavailable. They encode for multiple routes for producing the precursors for polyketide biosynthesis (acetyl-CoA, propionyl-CoA, and malonyl-CoA) including the capability to import, and degrade branched chain amino acids to these precursors. Finally, one of the *Ca*. Angelobacter genomes (*SRVP-Angelobacter-2*) from the vernal pool soil contained a Type IC CRISPR-Cas array and another (*SRVP-Angelobacter-3*) contained a Type IIID CRISPR-Cas system, indicating some degree of pressure from phage predation for these species.

### *Candidatus* Angelobacter species are transcriptionally active *in situ*

To track transcriptional activity of *Ca*. Angelobacter *in situ*, we flash-froze soil samples in the field to preserve for metatranscriptomic RNA extraction, extracting and sequencing 20 Gbp of RNA for 10 samples taken from the vernal pool study site in 2019. Mapping both DNA and RNA reads back to genomes obtained from the site, we were able to track both relative abundance (DNA) and relative transcriptional activity (RNA) for microbes of interest. Across organisms, we found that microbial genomes that recruited more DNA reads tended to also recruit more RNA reads, implying abundant microbes were often more active. Two *Ca*. Angelobacter species (*SRVP-2* and *SRVP-3*) were found to be more transcriptionally active than the mean or median bacterial/archaeal species genome obtained from the site (**Fig 4A**). Another genome, *SRVP-Angelobacter-1*, was not observed to have significant transcriptional activity in the RNA samples.

**Figure 4:**
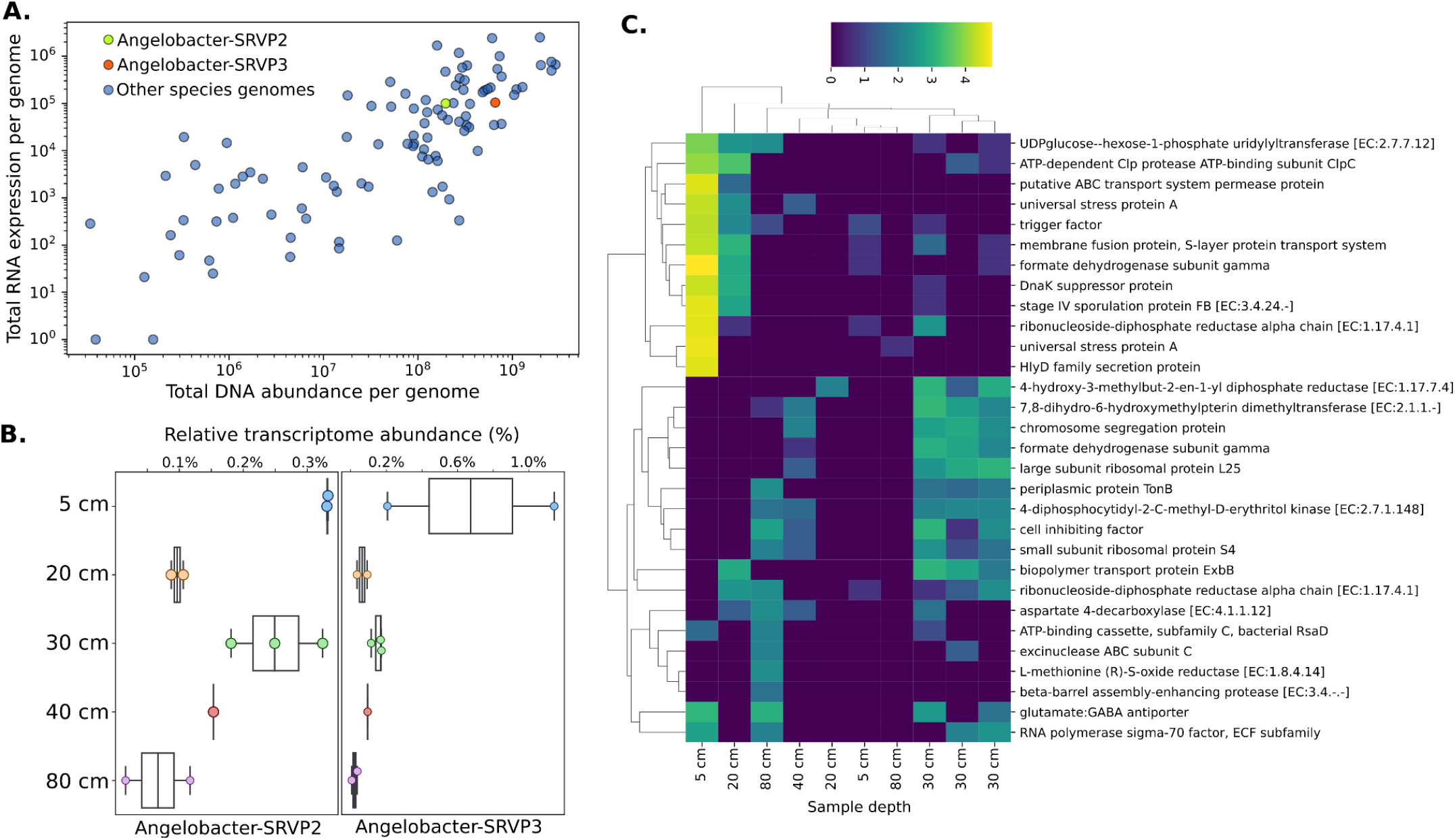
Metatranscriptomic activity of *Candidatus* Angelobacter species in situ. (**A**) Total metatranscriptomic (y-axis) and metagenomic (x-axis) reads mapping to two *Ca*. Angelobacter species, compared to all other microbes with genomes obtained from the vernal pool site. (**B**) Genome-wide transcriptional activity for two *Ca*. Angelobacter species, compared by sediment depth of sampling. (**C**) Transcriptomic counts of the most highly expressed Kegg Orthologs (KOs) at each sampling depth for the two *Ca*. Angelobacter species.

Overall, for the two active *Ca*. Angelobacter species (SRVP-2 and SRVP-3) we identified transcripts for 20% and 22% of their genes, respectively. In part, a low median level of detectable transcriptional activity can reflect the high diversity and low relative abundance of all bacteria in soils, but it could also be a consequence of large genomes that confer a high level of metabolic flexibility needed to adapt to changing conditions. We compared total expression for these two *Ca*. Angelobacter species by soil depth of the sample, and found that the species were most active at 5 cm depths, followed by 30 cm depths, and were least active in 80 cm deep soils (**Fig 4B**).

We next identified and annotated the 50 most abundant transcripts from either *Ca*. Angelobacter species in the dataset at both shallow (5-20 cm) and deeper depths (30-80 cm) (**Fig 4C**). Intriguingly, we observed a strongly unique transcriptomic profile in one sample taken from a 5 cm depth, in which several stress-related genes were highly expressed that remained largely unexpressed across the other samples. The highly expressed set of genes were clustered by their transcriptional abundances across samples, and samples did not cluster strongly by sample depth. However, three samples collected at a depth of 30 cm clustered together, with high expression for a set of genes that seemed to be involved in growth and general metabolism: ribosomal proteins, formate dehydrogenase, and a chromosomal segregation protein. We observed expression for about 5% of genes found in antiSMASH biosynthetic gene clusters, consistent with biosynthetic gene transcriptomic activity being on average lower than most cellular processes. A Type III PKS squalene-hopene cyclase, methyltransferases, and nonribosomal peptide synthetases were among the genes in BGCs with detectable levels of expression. Many of the genes with the highest levels of transcriptional activity in BGCs were transporter genes, including MacB-tripartite pumps and an AcrD multi drug efflux system. These data are consistent with *Ca*. Angelobacter transcriptional activity *in situ*, with activity highest at shallower soil depths, and possibly strong changes in transcriptional regulation between samples, similar to the kind previously reported in the previous metatranscriptomic experiment performed on soils with the originally reported strain.

## Discussion

This research substantially expanded upon a prior finding that the genus *Ca*. Angelobacter is genomically enriched in biosynthetic gene content. While metagenome-derived genomes from this lineage have been reported before, here we substantially expand sampling and provide evidence that this is a general feature of species in this genus. The results drew upon data from a very wide diversity of soil types, ranging from relatively dry soils that experience a Mediterranean climate (Angelo Reserve, with little or no rainfall for ~ 5 months per year) to permanently wet soils of a vernal pool. *Ca*. Angelobacter genomes were also identified in publicly available metagenomic datasets from agricultural soil, forest soils, and soil amended with pyrogenic organic matter. The relatively large gene inventories of these Acidobacteria may confer the metabolic flexibility needed to proliferate over a range of soil conditions. Given their ability to respire and ferment complex organic carbon compounds and their abundance and activity in saturated, organic carbon rich soil *Ca*. Angelobacter may also contribute substantially to carbon compound turnover.

Intriguingly, these specialized metabolite producers were much more abundant in the soils from this study than bacteria commonly cultivated from soil and known for their ability to produce specialized metabolites. For example, we only recovered marker genes from *Streptomyces* or *Pseudomonas* at low abundances in this study. We also demonstrate that *Ca*. Angelobacter genomes reconstructed in this study encode for as many, and sometimes more, biosynthetic genes than the originally reported genome. Further, the majority of these BGCs are unrelated, indicating substantial biosynthetic gene diversity within the genus. While functional prediction of BGCs is challenging, we identified *Ca*. Angelobacter BGCs containing MacB tripartite efflux pumps, yet none with siderophore-specific TonB-dependent receptors. This could indicate that many of these BGCs are involved in interbacterial competition rather than iron acquisition. Consequently, isolation of these bacteria and chemical characterization of their products may represent a priority for discovery of pharmaceutically important natural products.

While genome-resolved metagenomics presents its own biases as a window into soil microbial community composition - particularly sequencing bias against high GC% genes, poor DNA extraction from spores, and difficult assembly of high strain complexity species - the findings of this study imply that as yet uncultivated Acidobacteria may play an outsized role in chemical ecology in soil ecosystems.

## Code and data availability

The Acidobacteria genomes and raw sequencing reads for this study will be made available under NCBI BioProject number PRJNA728365. Genomes and biosynthetic gene clusters are also made available at https://figshare.com/projects/A_widely_distributed_genus_of_soil_Acidobacteria_genomically_enriched_in_biosynthetic_gene_clusters/113286.

## Acknowledgements

We gratefully acknowledge the Innovative Genomics Institute and the Chan Zuckerburg Biohub for sequencing resources and funding. We also gratefully acknowledge Dr. Thea Whitman for sharing metagenomic data for some of the findings in this manuscript.

## Competing Interests

The authors declare that they have no conflict of interest.

---

**Figure S1:**
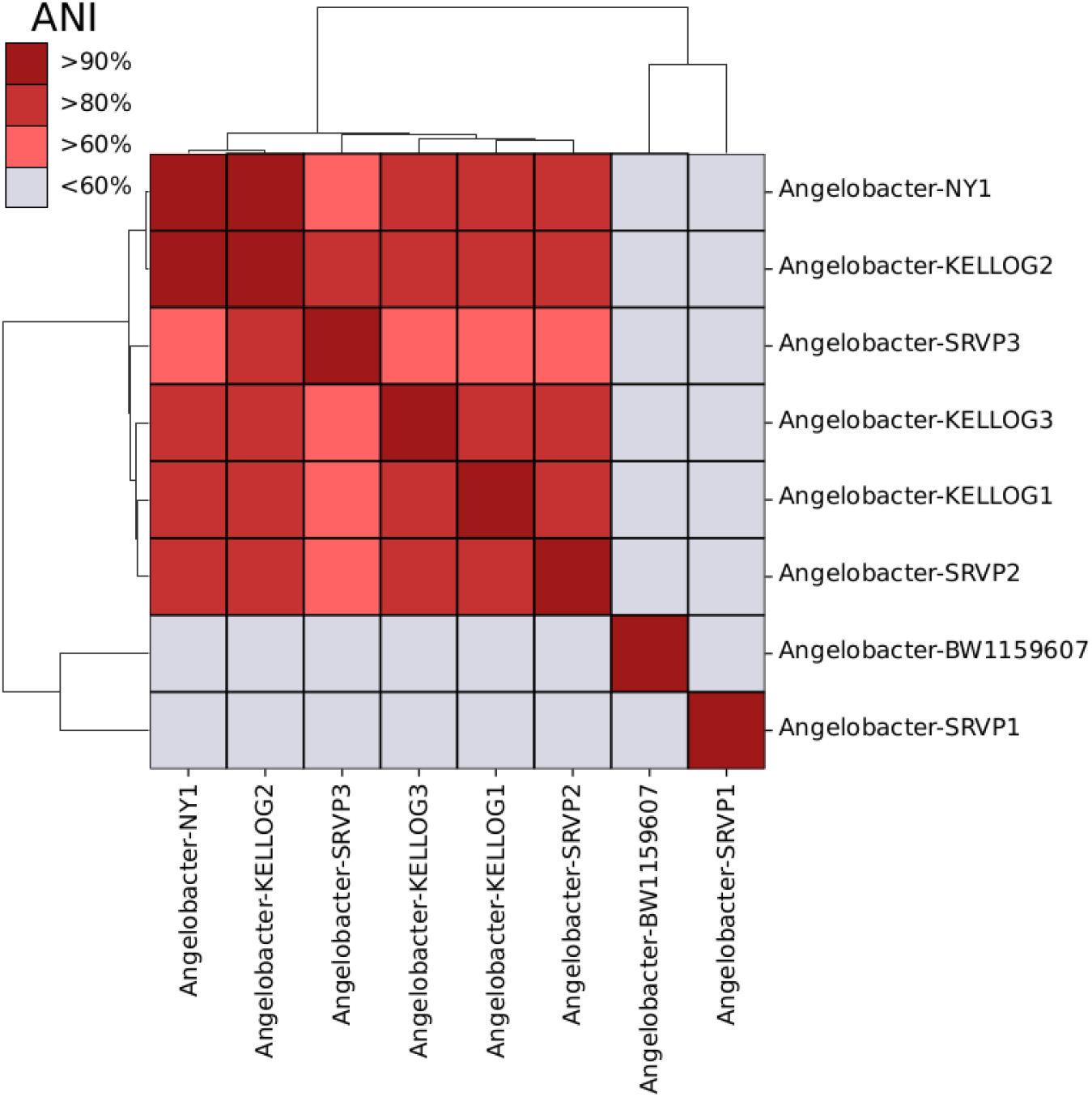
Genomic sequence identity between *Ca*. Angelobacter species obtained during this study.

**Figure S2:**
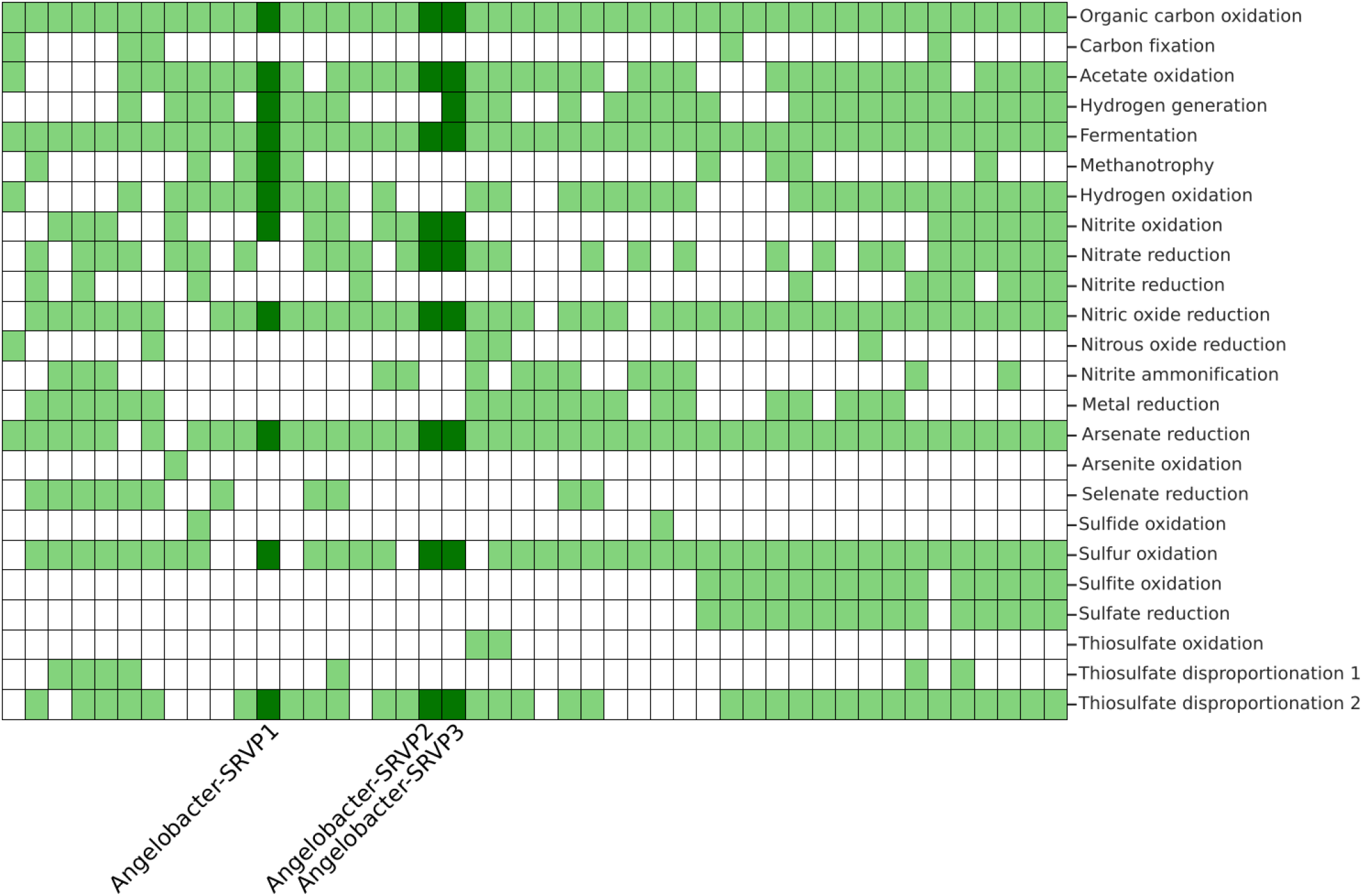
Genomic potential for generalized biogeochemical pathways in Acidobacterial genomes from the vernal pool (1 genome per column). A pathway is marked as present if the core kegg orthologs encoding that pathway are identified in each genome with METABOLIC.

## Notes

### Competing Interest Statement

The authors have declared no competing interest.

